# A subset of nucleus accumbens neurons receiving dense and functional prelimbic cortical input are required for cocaine seeking

**DOI:** 10.1101/2021.09.03.458762

**Authors:** Benjamin M. Siemsen, Sarah M. Barry, Kelsey Vollmer, Lisa M. Green, Ashley G. Brock, Annaka M. Westphal, Raven A. King, Derek M. DeVries, James M. Otis, Christopher W. Cowan, Michael D. Scofield

**Affiliations:** Department of Anesthesiology and Perioperative Medicine, Medical University of South Carolina, Charleston, SC; Department of Anatomy and Neurobiology, University of Maryland School of Medicine, Baltimore, MD; Department of Neuroscience, Medical University of South Carolina, Charleston, SC

**Keywords:** Cocaine, Prelimbic, Nucleus accumbens, Sucrose, Glutamate, Astrocytes

## Abstract

**Background:** Prelimbic cortical projections to the nucleus accumbens core are critical for cue-induced cocaine seeking, but the identity of the accumbens neuron(s) targeted by this projection, and the transient neuroadaptations contributing to relapse within these cells, remain unknown.

**Methods:** Male Sprague-Dawley rats underwent cocaine or sucrose self-administration, extinction, and cue-induced reinstatement. Pathway-specific chemogenetics, patch-clamp electrophysiology, *in vivo* electrochemistry, and high-resolution confocal microscopy were used to identify and characterize a small population of nucleus accumbens core neurons that receive dense prelimbic cortical input to determine their role in regulating cue-induced cocaine and natural reward seeking.

**Results:** Chemogenetic inhibition of prelimbic cortical projections to the nucleus accumbens core suppressed cue-induced cocaine relapse and normalized real-time cue-evoked increases in accumbens glutamate release to that of sucrose seeking animals. Furthermore, chemogenetic inhibition of the population of nucleus accumbens core neurons receiving the densest prelimbic cortical input suppressed cocaine, but not sucrose seeking. These neurons also underwent morphological plasticity during the peak of cocaine seeking in the form of dendritic spine expansion and increased ensheathment by astroglial processes at large spines.

**Conclusions:** We identified and characterized a unique subpopulation of nucleus accumbens neurons that receive dense prelimbic cortical input. The functional specificity of this subpopulation is underscored by their ability to mediate cue-induced cocaine relapse, but not sucrose seeking. This subset of cells represents a novel target for addiction therapeutics revealed by anterograde targeting to interrogate functional circuits imbedded within a known network.

## Introduction

Relapse to cocaine use can be precipitated by cues or contexts predicting cocaine (1) which activate key cortical and limbic regions involved in craving (2). Pre-clinical models of cue-induced cocaine craving rely upon the use of self-administration (SA), abstinence (with or without extinction training), and re-exposure to cocaine-paired cues or contexts, which is crucial for interrogating neuronal circuits and cell types mediating cocaine relapse (3).

Cue-mediated activation of prelimbic (PrL) cortical glutamatergic neurons, particularly their projection to the nucleus accumbens core (NAcore), is required for reinstatement of cocaine seeking after extinction (4, 5). Within the NAcore, the homeostatic regulation of glutamate is disrupted following extinction of cocaine self-administration (6) due to dysfunction in astrocyte-mediated glutamate clearance (7), among other adaptations (8-10). Together, cocaine-induced alterations in neural function yield enhanced glutamate release in the NAcore when animals are exposed to cocaine cues, driving drug seeking (11, 12). Accordingly, pharmacological inhibition of the PrL cortex (13) or optogenetic inhibition of PrL terminals in the NAcore (14, 15) suppresses cue-induced cocaine seeking, without affecting seeking for natural rewards (16). Downstream from the PrL cortex, NAcore MSNs integrate inputs from multiple regions to guide motivated behavior (17). PrL inputs represent a major driver of NAcore neurons (18) and neural processes within the NAcore that guide motivated behavior are known to be disrupted by cocaine (see (19-21) for reviews). While electrical stimulation of the PFC evokes excitatory post-synaptic potentials (EPSPs) in the vast majority of NAcore neurons, action potential firing is observed in less than half of NAcore neurons (18, 22). Additionally, when relatively weak PFC stimulation is employed,∼20% of NAcore neurons exhibit spike firing, suggesting that a subpopulation of NAcore neurons receives the majority of PrL inputs (18) and that there is heterogeneity of functional innervation of NAcore MSNs by PrL neurons. Given the importance of PrL afferents in driving cocaine seeking,the NAcore neurons receiving the majority of PrL inputs are a particularly relevant population for understanding relapse vulnerability.

Cue-induced reinstatement of several classes of drugs, including cocaine, elicits a transient increase in synaptic strength in NAcore medium spiny neurons (MSNs) (23-25). During the peak of cocaine seeking, NAcore MSNs undergo PrL-dependent morphological and synaptic plasticity (enhanced dendritic spine head diameter and AMPA/NMDA ratio) – events that are required for cue-induced cocaine seeking (23). Moreover, following cocaine or heroin SA, astrocyte processes exhibit reduced contact with bulk NAcore synapses (*i*.*e*. Synapsin-I^+^ synapses), which contributes to heightened cue-induced glutamatergic signaling. During cued heroin seeking, astrocyte processes “re-associate” with NAcore synapses, presumably to limit cue-induced glutamate signaling, dampening heroin seeking (26). However, the specific MSN population whereby increased astrocyte association occurs, whether this occurs on functionally relevant synapses, and whether this holds true for cue-induced cocaine seeking, is unknown. This is important given that astrocyte regulation of synaptic transmission is highly specific and that astrocytic processes can interact and regulate both excitatory and inhibitory synapses in response to changes in overall network activity (27-29).

In this study, we systematically evaluated the contribution of PrL neurons projecting to the NAcore (PrL^NAcore^) in cue-induced cocaine and sucrose seeking using an intersectional chemogenetic viral vector approach, while simultaneously measuring glutamate release in the NAcore during seeking. We then identified a novel sub-population of NAcore neurons that receive the most dense and active PrL cortical innervation (NAcore^PrL^) using an anterograde, transsynaptic AAV1-Cre vector in the PrL and Cre-dependent constructs in the NAcore. We used this combinatorial viral vector approach to examine a) the role of NAcore^PrL^ neurons in cocaine and sucrose seeking and b) the regulation of NAcore^PrL^ synapses and their ensheathing astrocytic processes in cue-induced cocaine seeking.

## Materials and methods

### Animal subjects and surgery

Male Sprague-Dawley rats (*N*=82) were used for all experiments. All animal use protocols were approved by the Medical University of South Carolina and were performed according to the National Institutes of Health Guide for the Care and Use of Laboratory Animals (8^th^ ed., 2011). Viral vector information can be found in Table 1. Detailed methods can be found in the supplemental information.

**Table 1.** Virus, source, titer, injection location, and purpose for all viral vectors used.

| **Virus** | **Source** | **Titer**<br>**(vg/ml)** | **Injection**<br>**site** | **Purpose** |
| --- | --- | --- | --- | --- |
| AAV1-CamKII-Cre-0.4-SV40 | Addgene | 1.2-1.9 X 10<sup>13</sup> | PrL | Transsynaptic Cre expression |
| AAV5-GfaABC1D-Lck-GFP | University of North Carolina Vector Core | 0.55 X 10<sup>13</sup> | NAcore | GFP expression in astrocyte membrane processes |
| AAV2-hSyn-DIO-hM4Di | Addgene | 2.5 X 10<sup>13</sup> | PrL;<br>NAcore | Cre-dependent hM4Di expression |
| AAV2-hSyn-DIO-mCherry | Addgene | 1.6 X 10<sup>13</sup> | PrL;<br>NAcore | Cre-dependent mCherry expression |
| AAV1-CAG-Flex-Ruby2sm-Flag-WPRE-SV40 | Addgene | 1.1 X 10<sup>13</sup> | NAcore | Cre-dependent expression of Flag-tagged smFP |
| AAV2-EF1a-DIO-hChR2(H134R)-EYFP | University of North Carolina Vector Core | 1.6 X 10<sup>12</sup> | PrL | Cre-dependent Channel Rhodopsin expression |
| AAVrg-hSyn-Cre-WPRE-hGH | Addgene | 2.1 X 10<sup>13</sup> | NAcore | Expression of Cre in neurons projecting to NAcore |

### Drugs

Cocaine hydrochloride (NIDA, Research Triangle Park NC; 200 µg/50 µl bolus) was dissolved in sterile saline for SA experiments. Clozapine-N-Oxide (CNO) was obtained through the National Institute of Mental Health Chemical Synthesis Program. CNO was dissolved in 5% DMSO in sterile saline and was administered at a dose of 5 mg/kg (i.p.) 30 minutes before testing.

### Cocaine and sucrose SA

Active lever presses (ALP), inactive lever presses (ILP) and infusions earned are shown within each figure for all SA experiments. Detailed methods can be found in the supplemental information.

### Extinction and cue-induced reinstatement

During daily extinction sessions (2 hours/day) ALPs had no programmed consequence. Animals underwent extinction sessions until they met criterion (average ≤ 25 ALPs over the last 2 days). Rats then underwent a cue-induced reinstatement test whereby ALPs elicited the light and tone cue-complex, but no cocaine or sucrose delivery. Yoked saline animals were sacrificed 24 hours after the final extinction session without undergoing reinstatement. At the beginning of each reinstatement session, a single noncontingent cue was presented. Following the 2-hour reinstatement session, a subset of rats were perfused 15 minutes after the session for Fos immunohistochemistry (see below). Virus expression in both the PrL cortex and NAcore were mapped according to the atlas of Paxinos and Watson.

### Perfusions and immunohistochemistry

Antisera information can be found in the key resource table. Detailed methods can be found in the supplemental information.

### Patch-clamp electrophysiology

Rats were anesthetized with isoflurane and rapidly decapitated for brain extraction. Brains were sectioned (300*μ*m) using a vibratome (Leica VT 1200) in ice-cold, oxygenated sucrose cutting solution containing (in mM): 225 sucrose, 119 NaCl, 1.0 NaH_2_PO_4_, 4.9 MgCl_2_, 0.1 CaCl_2_, 26.2 NaHCO_3_, 1.25 glucose, ∼305 mOsm. Brain slices recovered for 30 mins in warm (32°C) aCSF containing (in mM): 119 NaCl, 2.5 KCl, 1.0 NaH_2_PO_4_, 1.3 MgCl, 2.5 CaCl_2_, 26.2 NaHCO_3_, 15 glucose, ∼305 mOsm. During recording, slices were perfused with room-temperature aCSF (1 mL/min). NAcore cells were visualized using differential interference contrast (DIC) through a 40x liquid-immersion objective mounted on an upright light microscope (Olympus, BX-RFA). NAcore MSNs expressing mCherry were visualized using an integrated green LED (545nm; < 1mW). Whole-cell recordings were obtained using borosilicate pipettes (3-7 M*Ω*) back-filled with Cesium methanesulfonate, which contained (in mM): 117 Cs methanesulfonic acid, 20 HEPES, 10 EGTA, 2 MgCl_2_, 2 ATP, 0.2 GTP (pH 7.35, mOsm 280).

Voltage-clamp recordings were obtained from mCherry^+^ and neighboring mCherry^-^ NAcore MSNs to characterize inputs from PrL. During recordings, NAcore MSNs were held at - 70mV and PrL axons containing AAV2-EF1a-DIO-hChR2-EYFP were activated through a 10ms pulse of a blue LED (470nm; 1 mW) delivered every 5s. The resulting optogenetically-evoked EPSCs were recorded and peak amplitudes were analyzed using Clampfit (v10).

### *In Vivo* electrochemistry

*In vivo* electrochemical detection and quantification of glutamate levels were performed as previously described (12). Following SA, surgery, abstinence, and extinction, Glutamate oxidase (GluOx)-coated electrochemical electrodes were implanted (Pinnacle Technology, Lawrence, KS) and connected to a wireless potentiostat housed within a 3D-printed enclosure; data was transmitted from the potentiostat via Bluetooth. Electrodes were calibrated and implanted as described previously (12). Following implantation animals were returned to the colony overnight. The next morning, a ∼1-hour baseline recording began; animals undergoing cocaine seeking were injected with CNO (5 mg/kg, i.p.) 30 minutes prior to the reinstatement session. Animals undergoing sucrose seeking did not receive CNO. 1Hz measurements of glutamate-dependent currents were expressed relative to a 15-minute pre-session baseline, then converted to glutamate concentrations (in nM) based off the *in vitro* calibration. While behavioral testing lasted two hours, recordings were stopped after the first hour. Animals were perfused following testing. Virus expression and probe placement were mapped according to the atlas of Paxinos and Watson.

### Microscopy

Detailed methods regarding virus vector mapping, cell counting, and Fos imaging and analyses can be found in the supplemental information.

#### Dendritic spine and astrocyte membrane imaging

Eighty-micron (μm) coronal sections were immunohistochemically processed for Flag (to label neurons) and GFP (to label astrocytes) and imaged using a Leica SP8 laser-scanning confocal microscope. NAcore neurons receiving PrL input and surrounding astrocytes were imaged with an OPSL 552nm and an Argon 488nm laser line, respectively. Dendrites were selected for imaging based off the following criteria: 1) relative isolation from interfering dendrites, 2) location past the second branch point from the soma, 3) traceability back to a soma of origin, and 4) location within a field of labeled astrocytes. Images were acquired using a 63X oil-immersion objective (1.4 N.A.) with a frame size of 1024×512 pixels, a step size of 0.1 µm, 4.1X digital zoom, and a 0.8AU pinhole. Laser power and gain were optimized then held relatively constant only adjusting to maintain voxel saturation consistency between images. Images were deconvolved using Huygens software (Scientific Volume Imaging, Hilversum, Netherlands) prior to analyses.

### Imaris analyses

#### Dendritic spine morphometrics and astrocyte association at dendritic spine heads

Deconvolved Z-stacks were imported to Imaris. Dendritic spine analyses were performed as previously described (30). Briefly, the filament tool was used to trace the dendrite shaft and spines. The average diameter (d_H_, µm) of spine heads on each dendrite, as well as the dendritic spine density (number of spines per µm of dendrite) were exported.

To analyze astrocyte association with spines (Figure S1), the filament analysis extension was used to model dendritic spine heads as spheres, which were converted to rendered surfaces. Next, the dilate surface extension was used to expand each dendritic spine head surface by 300 nm, producing a hollow expanded sphere for each spine head. Next, an automatic threshold was used to render the volume of astrocyte membrane within each hollow sphere ROI. The physical volume of astrocyte signal within each hollow sphere ROI surrounding the spine head was normalized to the corresponding volume of each hollow sphere, yielding a % association for each spine head. This allows equivalent analyses of astrocyte association with dendritic spine heads, independent of the size of the spine head. Average GFP intensity within the ROI, % of spines along the dendrite with 0 association, the average % association at spine heads, as well as the % association at each individual spine head on the dendrite were assessed.

### Statistical analyses

All statistical analyses were performed with GraphPad Prism (v9) software. When comparing groups across time or bins a mixed-effects model (for missing values) or mixed-model two-way ANOVA with virus or treatment as a between subject variable and session/test/bin as the within subject variable followed by Bonferroni-corrected multiple comparison test when an interaction or main effect was significant. Fos data were analyzed with a two-way ANOVA with virus (mCherry vs. hM4Di) and reward (cocaine vs. sucrose) as between subject variables followed by Tukey’s multiple comparison tests when an interaction was significant. Correlation analyses were performed using a two-tailed Pearson’s correlation. Electrochemical calibration data was analyzed with linear regression and changes in glutamate concentrations during reinstatement were analyzed using area under the curve (AUC) analyses. AUC’s were then compared across groups using a one-way ANOVA with a Dunnett’s multiple comparison test. Paired and unpaired observations were analyzed with a paired or un-paired, with the appropriate correction if needed, t-test, respectively. Significance was set at *p*<0.05 and data is expressed as the mean +/-the standard error of the mean. All non-significant behavioral data is presented in TableS1.

## Results

### Chemogenetic inhibition of PrL^NAcore^ neurons suppresses cue-induced reinstatement and associated glutamate release in the NAcore

To test whether PrL neurons projecting to the NAcore (PrL^NAcore^) are required for cue-induced reinstatement and increased glutamate release in the NAcore, we expressed inhibitory hM4Di-DREADDs in PrL^NAcore^ neurons using an intersectional chemogenetic approach (Figure 1A). Animals underwent cocaine or sucrose SA, followed by one week of home cage abstinence, then extinction training. Following cocaine or sucrose SA, 7-day home-cage abstinence, and extinction training (Figure 1C and Supplemental Fig S2). Animals were then administered CNO 30 minutes prior to the presentation of cocaine-paired cues. Cocaine seeking was then measured as the number of non-reinforced presses of the lever formerly paired with cocaine (*i*.*e*. “active” lever presses or ALPs). Comparing the last day of extinction, cue presentation to DIO-mCherry control animals produced a significant increase in drug seeking on the active lever (two-way mixed-model ANOVA, significant group by test interaction, F(1,21)=11.73, *p*=0.003, Figure 1D), which was suppressed in the DIO-hM4Di rats compared to mCherry controls (*p*<0.0001). When ALPs during reinstatement were summated in 15 minute bins, there was a main effect of group (F(1,21)=12.60, *p*=0.002, Figure 1E), indicating that ALPs were lower in hM4Di compared to mCherry rats regardless of the time bin. Representative PrL^NAcore^ hM4Di expression is shown in Figure 1F and a compiled heat map for hM4Di rats is shown in Figure 1G.

**Figure 1.**
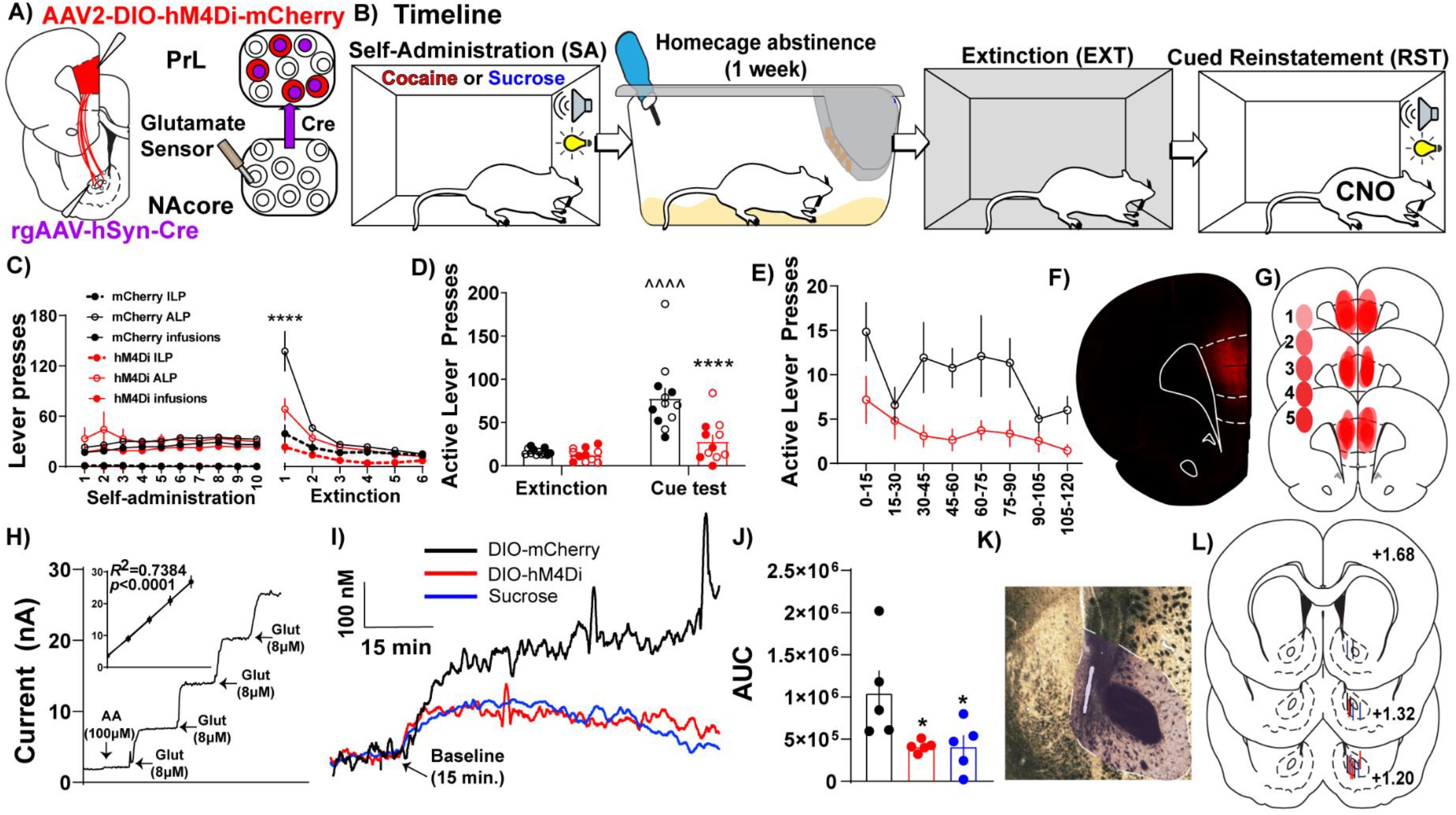
PrL^NAcore^ activity is required for cue-induced cocaine seeking after abstinence and extinction as well as cue-induced glutamate release in the NAcore. **A)** Surgery schematic for chemogenetic inhibition of PrL^NAcore^ neurons with simultaneous recording of glutamate release in the NAcore. **B)** Behavioral timeline. **C)** Cocaine SA and extinction inactive lever presses (ILP), active lever presses (ALP), and infusions earned in mCherry and hM4Di treatment groups. One hM4Di rat was removed for lack of extinction. Final group sizes: hM4Di (*n*=11) and mCherry (*n*=12) animals. **D)** Active lever presses during the last two days of extinction compared to cue-induced reinstatement in mCherry and hM4Di rats.^^^^*p*<0.0001 compared to extinction, \*\*\*\**p*<0.0001 compared to mCherry. Filled circles indicate animals that underwent *in vivo* glutamate recordings. **E)** Active lever presses during cue-induced reinstatement summated in 15-minute bins. **F)** Representative micrograph of PrL^NAcore^ hM4Di virus expression. **G)** Heat map of virus expression in hM4Di rats. Cooler shades indicate lower number of overlapping virus expression, and warmer shades indicate greater overlap. Numbers indicate the number of animals showing overlap. **H)** Representative Glu-Ox electrode *in vitro* calibration. Inset shows linear regression for all electrodes used (*n*=15). **I)** Average of 1Hz recordings of glutamate currents in the NAcore. **J)** AUC for all animals (*n*=5/group) \**p*<0.05 compared to mCherry. **K)** Representative micrograph of electrode placement in the NAcore. **L)** Compiled electrode placements; electrodes are color coded based off of treatment.

In a subset of the cocaine SA animals, we also examined the influence of PrL^NAcore^ neurons on cue-induced glutamate release in the NAcore. Extracellular glutamate levels were detected using a calibrated GluOx-coated electrode implanted in the NAcore (Figures 1A, 1H) and cued glutamate release was measured in cocaine (mCherry and hM4Di) and sucrose reinstating rats (Figure 1I). When analyzing the area under the curve (AUC) of 1Hz measurements of glutamate release in the three groups, we detected a main effect of cue on NAcore glutamate levels (one-way ANOVA, main effect of treatment, F(2,12)=4.55, *p*=0.034, Figure 1J). Compared to PrL^NAcore^ DIO-mCherry controls, the PrL^NAcore^ DIO-hM4Di animals and animals reinstating to sucrose-paired cues both showed a significant reduction in cue-induced glutamate release in the NAcore (*p*=0.04). Of note, the inhibition of PrL^NAcore^ projections reduced the cue-induced glutamate release to levels similar to those produced in animals reinstating to sucrose-paired cues, suggesting that PrL^NAcore^ represents a majority of the glutamatergic inputs promoting cue-triggered cocaine seeking.

### AAV1-CamKIIa-Cre injections in the PrL cortex transduce a subpopulation of NAcore neurons receiving the densest PrL cortical input

Like other AAV serotypes, AAV1 viral particles transduce neurons in the primary injection site. However, one unique property of AAV1 vectors is that a portion of viral particles are trafficked anterogradely down axons then released via vesicle fusion at axon terminals. This allows for monosynaptic transfer of viral particles and transgene expression in downstream neurons receiving synaptic contacts (31, 32). Importantly, AAV1-mediated transsynaptic transduction is strongest in neurons that receive numerous functional inputs from cells at the primary injection site. As such, we took advantage of this characteristic of AAV1 vectors to investigate NAcore neurons receiving dense and functional PrL inputs (NAcore^PrL^). First, we determined what percentage of NAcore neurons are transduced via AAV1 transsynaptic transduction by infusing AAV1-CamKII-Cre in the PrL cortex and AAV1-CAG-Flex-Ruby2sm-Flag (Flex-smFP) or AAV2-hSyn-DIO-hM4Di-mCherry in the NAcore (Figure 2A-C). Within the field of viral transduction in the NAcore, ∼10% of the neurons were labeled (Figure 2D). Further analysis of NAcore^PrL^ neurons revealed that a large fraction of NAcore^PrL^ neurons exhibited reactivity for pre-pro Enkephalin (ppENK), a marker for D2 MSNs (33), whereas a smaller subset exhibited reactivity for various interneuron markers (Figure 2E). Interestingly, even though a minority of NAcore^PrL^ neurons were identified as parvalbumin interneurons (PV), we found ∼60% of total NAcore PV neurons were strongly innervated by the PrL (Figure 2F).

**Figure 2.**
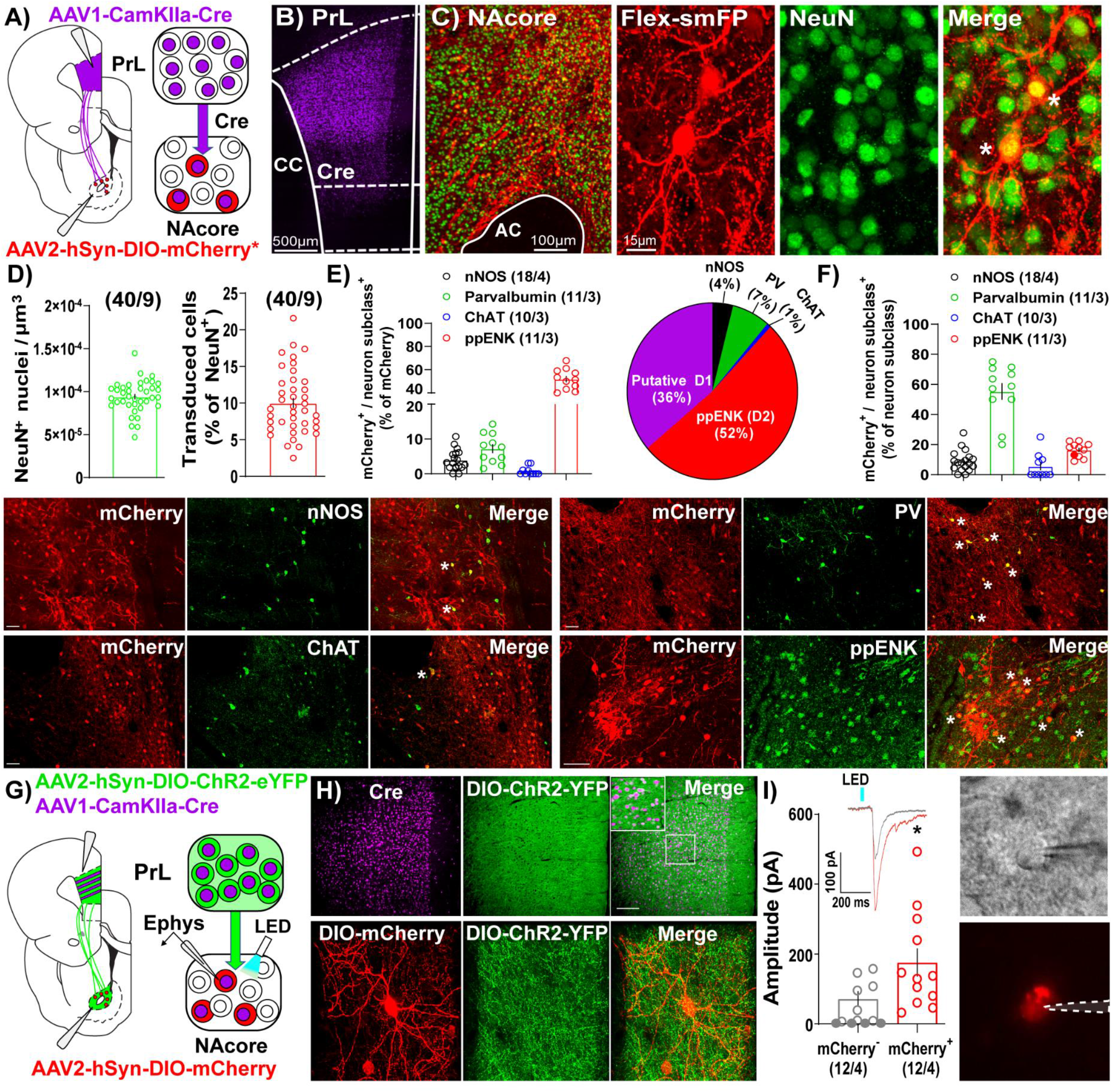
Immunohistochemical and electrophysiological characterization of NAcore^PrL^ neurons. **A)** surgery schematic (left) for labeling NAcore^PrL^ neurons for manipulation. Asterisk indicates use of DIO-hM4Di-mCherry or Flex-smFP for analyses. **B)** Representative confocal image of immunohistochemically-detected Cre (from AAV1-CamKIIa-Cre) in the first-order region (PrL cortex). **C)** Representative image of NeuN (green) and Flag-tagged Flex-smFP (red) in the NAcore. High magnification image is shown to the right. **D)** Left - NeuN^+^ nuclei normalized to dataset volume (µm^3^). Right - Percent of NeuN nuclei that were transduced by AAV1-CamKIIa-Cre in the PrL cortex and Cre-dependent vectors in the NAcore from three groups of rats, animals expressing Flex-smFP in NAcore^PrL^ neurons that underwent yoked-saline (11 images from 3 animals), 15 minutes of cocaine seeking (12 images from 3 animals), or animals undergoing two hours of cue-induced reinstatement with hM4Di expression in NAcore^PrL^ neurons (17 images from 3 animals). **E)** Left - Percent of NAcore^PrL^ neurons that were co-labeled by nNOS, Parvalbumin (PV), Choline acetyltransferase (ChAT), or pre-pro enkephalin (ppENK). Right - Pie chart summarizing data in E. **F)** Percent of each neuron subclass that was also mCherry^+^ (inverse of data in E). Representative images of mCherry/nNOS (top, left-scale bar=100µm), mCherry/ChAT (bottom, left-scale bar=100µm), mCherry/PV (top, right-scale bar=100µm), and mCherry/ppENK (bottom, right-scale bar=50µm). Tissue was processed from animals expressing DIO-hM4Di in NAcore^PrL^ neurons undergoing 2 hours of cue-induced reinstatement (*n*=3-4 animals). **G)** Schematic of slice electrophysiology during optogenetic stimulation of PrL terminals in the NAcore. **H)** Representative images of immunohistochemically-detected Cre in the PrL cortex (top, left), DIO-hChR2-EYFP (top, middle), and merge of the two signals (top, right) in the PrL cortex following an injection of AAV1-CamkIIa-Cre and AAV2-EF1a-DIO-hChR2-EYFP (*n*=4 rats). Scale bar=200 µm (top, main), 50 µm (inset). A representative DIO-mCherry-labeled NAcore^PrL^ neuron (bottom, left), axonal fibers from DIO-hChR2-EYFP in the NAcore (bottom, middle), and merge of the two signals (bottom, right) is shown below. Scale bar=30µm. **I)** Left - EPSC amplitude in mCherry^-^ and mCherry^+^ cells following optogenetic stimulation of PrL terminals in the NAcore. Inset shows representative traces from the two cell types. Right - Representative DIC (top) and fluorescent filtered (bottom) mCherry^+^ cell chosen for patching. \**p*<0.05 compared to mCherry^-^

In a separate cohort of drug-naïve rats, we co-injected AAV1-CamKIIa-Cre and AAV2-EF1a-DIO-hChR2-EYFP into the PrL cortex and AAV2-hSyn-DIO-mCherry in the NAcore and used a blue light-evoked EPSC protocol (34) to record EPSC amplitude in mCherry^+^ and neighboring mCherry^-^ neurons in the NAcore (Figure 2H-I). 58% of mCherry^-^ neurons either showed no response (33%), or a response <10 pA (25%), whereas 100% of mCherry^+^ cells responded. When removing mCherry^-^ cells that showed no response, mCherry^+^ cells showed a greater EPSC amplitude compared to mCherry^-^ cells (Welch’s-corrected t-test, t(16.34)=2.259, *p*=0.038, Figure 2J).

### Chemogenetic inhibition of NAcore^PrL^ neurons suppresses cue-induced reinstatement of cocaine, but not sucrose, seeking

To examine the contribution of Nacore^PrL^ neurons in reward-seeking behavior, we microinjected AAV1-CamKIIa-Cre into the PrL cortex and AAV2-hSyn-DIO-hM4Di-mCherry or AAV2-hSyn-DIO-mCherry into the NAcore (Figure 3A). Animals underwent cocaine or sucrose SA, extinction, and cue-induced reinstatement (Figure 3B). AAV1-CamKIIa-Cre injections were confined to the PrL cortex, and AAV2-DIO-hM4Di transduced a subpopulation of cells in the NAcore (Figure 3C-D). DIO-mCherry and DIO-hM4Di rats undergoing cocaine self-administration and extinction (Figure 3E) both reinstated to cocaine-paired cues (two-way mixed-model ANOVA, group by test interaction, F(1,21)=13.94, *p*=0.001; mCherry (*p*<0.0001), hM4Di (*p*=0.006), Figure 3F). Compared to mCherry controls, the chemogenetic inhibition of NAcore^PrL^ neurons reduced cue-induced reinstatement (*p*<0.0001), which was most pronounced during the first 30 minutes of the cue test when ALP data was binned into 15-minute time blocks (two-way RM ANOVA, virus by time point interaction, F(7,147)=3.743, *p*=0.0009, Figure 3G) and significant differences between mCherry and hM4Di were observed at 15 (*p*<0.0001) and 30 min (*p*<0.05) minutes. Interestingly, for DIO-mCherry and DIO-hM4Di rats undergoing sucrose self-administration and extinction (Figure 3H), there was a main effect of cued sucrose seeking (two-way mixed model ANOVA, F(1,14)=46.03, *p*<0.0001, Figure 3I), yet NAcore^PrL^ inhibition did not affect cue-induced reinstatement. There was also no difference in cued sucrose seeking with NAcore^PrL^ inhibition when ALP data was binned (two-way RM ANOVA, F(7,98)=0.82, *p*=0.573, Figure 3J), indicating that NAcore^PrL^ neurons are required for cued seeking for cocaine, but not natural rewards.

**Figure 3.**
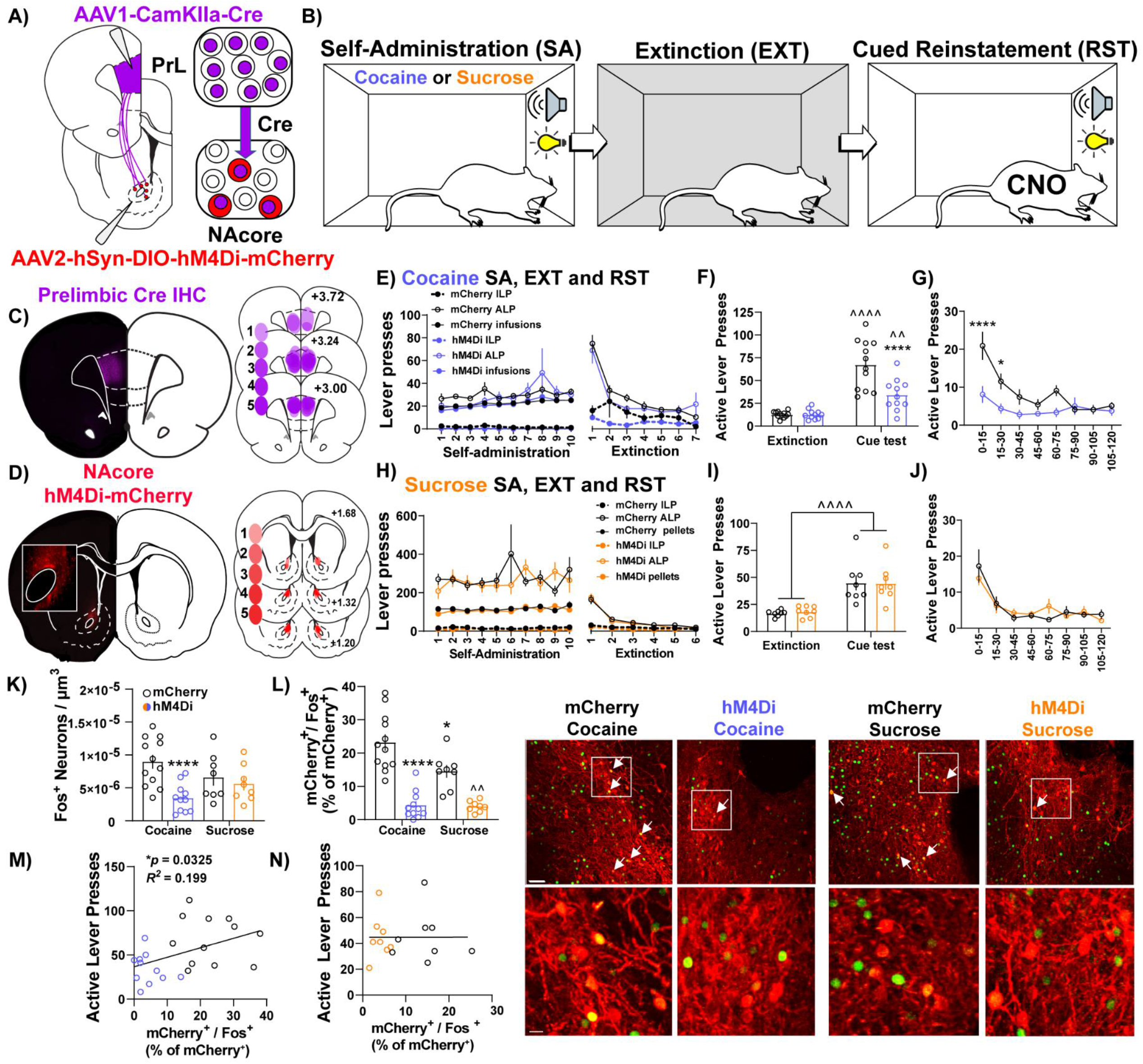
NAcore^PrL^ neuronal activity is a critical regulator of cue-induced cocaine, but not sucrose, seeking. **A)** Surgery schematic for expression of inhibitory DREADDs or mCherry control in NAcore^PrL^ neurons. Two hM4Di rats were removed from the analysis due to off-target virus expression in the NAcore. **B)** Behavioral timeline. Animals underwent either cocaine or sucrose SA, extinction, then cue-induced reinstatement (CNO 5 mg/kg 30 minutes before reinstatement). **C)** Representative immunohistochemically-detected Cre in the PrL cortex (left) and a heatmap of viral spread in the PrL cortex of hM4Di-expressing rats. Cooler shades indicate lower number of overlapping virus expression, and warmer shades indicate greater overlap. Numbers indicate the number of animals showing overlap. **D)** Representative DIO-hM4Di-mCherry expression in the NAcore (left) and a heatmap of viral spread in the NAcore of hM4Di-expressing rats. **E)** Cocaine SA and extinction inactive lever presses (ILP), active lever presses (ALP), and infusions earned in mCherry (*n*=12) and hM4Di (*n*=11) groups. **F)** Average last 2 days of extinction ALP (left) and ALP during cue-induced reinstatement (right) in two treatment groups undergoing cue-induced reinstatement of cocaine seeking. ^^*p*<0.01, ^^^^*p*<0.0001 compared to extinction, \*\*\*\**p*<0.0001 compared to mCherry. **G)** ALP during reinstatement in 15-minute bins.\**p*<0.05, \*\*\*\**p*<0.0001 compared to mCherry. **H)** Sucrose SA (*n*=8 per group) and extinction ILP, ALP, and pellets earned in the two treatment groups. **I)** Average last 2 days of extinction ALP (left) and ALP during cue-induced reinstatement (right) in two treatment groups undergoing cue-induced reinstatement of sucrose seeking. **J)** ALP during reinstatement in 15-minute bins for animals undergoing cue-induced reinstatement of sucrose seeking. **K-L)** Fos^+^ neurons (K - normalized to dataset volume, µm^3^) and percentage of NAcore^PrL^ neurons that were Fos^+^ (L) in the four groups undergoing cue-induced reinstatement (mCherry-cocaine, hM4Di-cocaine, mCherry-sucrose, hM4Di-sucrose). \**p*<0.05, \*\*\*\**p*<0.0001 compared to mCherry-cocaine.^^*p*<0.01 compared to mCherry-sucrose. **M-N)** Correlation of the percentage of NAcore^PrL^ neurons that were Fos^+^ with active lever presses during reinstatement in cocaine (M) and sucrose(N) reinstating rats. Representative confocal images of Fos (green) and mCherry (Red) in the four treatment groups. Bottom row are insets of boxed region in top row. Arrows indicate mCherry^+^/Fos^+^ neurons. Scale bars=30µm (top row) and 20µm (bottom row).

Next, we used Fos immunoreactivity to examine neuronal activation in the NAcore following cued reinstatement of cocaine or sucrose seeking. Compared to yoked-saline controls sacrificed 24 hours after the last extinction session, cued seeking in virus control animals (mCherry-Cocaine and mCherry-sucrose) produced a significant increase in Fos staining in the NAcore, which included NAcore^PrL^ neurons (Figure S3). When also including cue-induced Fos staining in the hM4Di-Cocaine and hM4Di-sucrose groups, a significant interaction between reward and virus was observed (two-way ANOVA, F(1,35)=5.178, *p*=0.029, Figure 3K). Compared to mCherry-Cocaine animals, only hM4Di-Cocaine animals showed reduced Fos in the NAcore overall (*p*=0.0007). When limiting the analysis to NAcore^PrL^ neurons, a two-way ANOVA revealed a significant reward by virus interaction (F(1,35)=4.746, *p*=0.036, Figure 3L). Compared to mCherry-Cocaine rats, all other groups showed reduced Fos expression in NAcore^PrL^ neurons (*p*’s<0.05). As expected, hM4Di-Sucrose rats showed reduced Fos expression in NAcore^PrL^ neurons compared to mCherry-Sucrose (*p*=0.005). Interestingly, ALPs during reinstatement positively correlated with Fos activation in NAcore^PrL^ neurons in cocaine (*r*=0.447, *p*=0.033, Figure 3M), but not sucrose animals (*r*=0.002, *p*=0.992, Figure 3N).

### NAcore^PrL^ MSNs undergo transient structural plasticity and increased astrocyte association at dendritic spines during reinstatement

Given NAcore^PrL^ neurons are essential for cued drug seeking, we next asked whether NAcore^PrL^ dendritic spine heads and surrounding perisynaptic astrocyte processes (PAPs) undergo morphological plasticity during cued cocaine seeking. To label NAcore^PrL^ neurons and surrounding astrocytes, we microinjected AAV1-CamKIIa-Cre in the PrL cortex and a mix of AAV1-CAG-Flex-Ruby2sm-Flag to label dendritic spines and AAV5-GfaABC1D-Lck-GFP to label astrocyte PAPs in the NAcore (Figure 4A). Following cocaine SA and extinction training, animals were sacrificed 15 minutes into cue-induced reinstatement (cocaine), and these were compared to yoked-saline control animals sacrificed 24 hours after extinction (Figure 4B). There was no difference in the percent of transduced neurons in the NAcore between cocaine and yoked saline rats (t(21)=0.794, *p*=0.436, Figure 2D), or in the percentage of NAcore^PrL^ neurons that were ppENK^+^ (Figure S4). Akin to prior experiments, CamKIIa-Cre was confined to the PrL cortex and Flex-Ruby2sm-Flag provided sparse, fully labeled MSNs for spine analyses (Figure 4C-D). Following cocaine SA and extinction training (Figure 4E), we observed that cued reinstatement significantly increased dendritic spine head diameter (d_H_) of NAcore^PrL^ spines (paired t-test, t(115)=5.780, *p*<0.0001, Figure 4F). This increase in d_H_ was associated with a rightward shift in dendritic spine d_H_ when spine d_H_ was binned to generate a frequency distribution, indicating a general increase in the size of spine heads (two-way mixed model ANOVA, group by bin interaction, F(5,575)=13.22, *p*<0.0001, Figure 4G), with no change in spine density (t(115)=0.905, *p*=0.367, Figure 4H-I).

**Figure 4.**
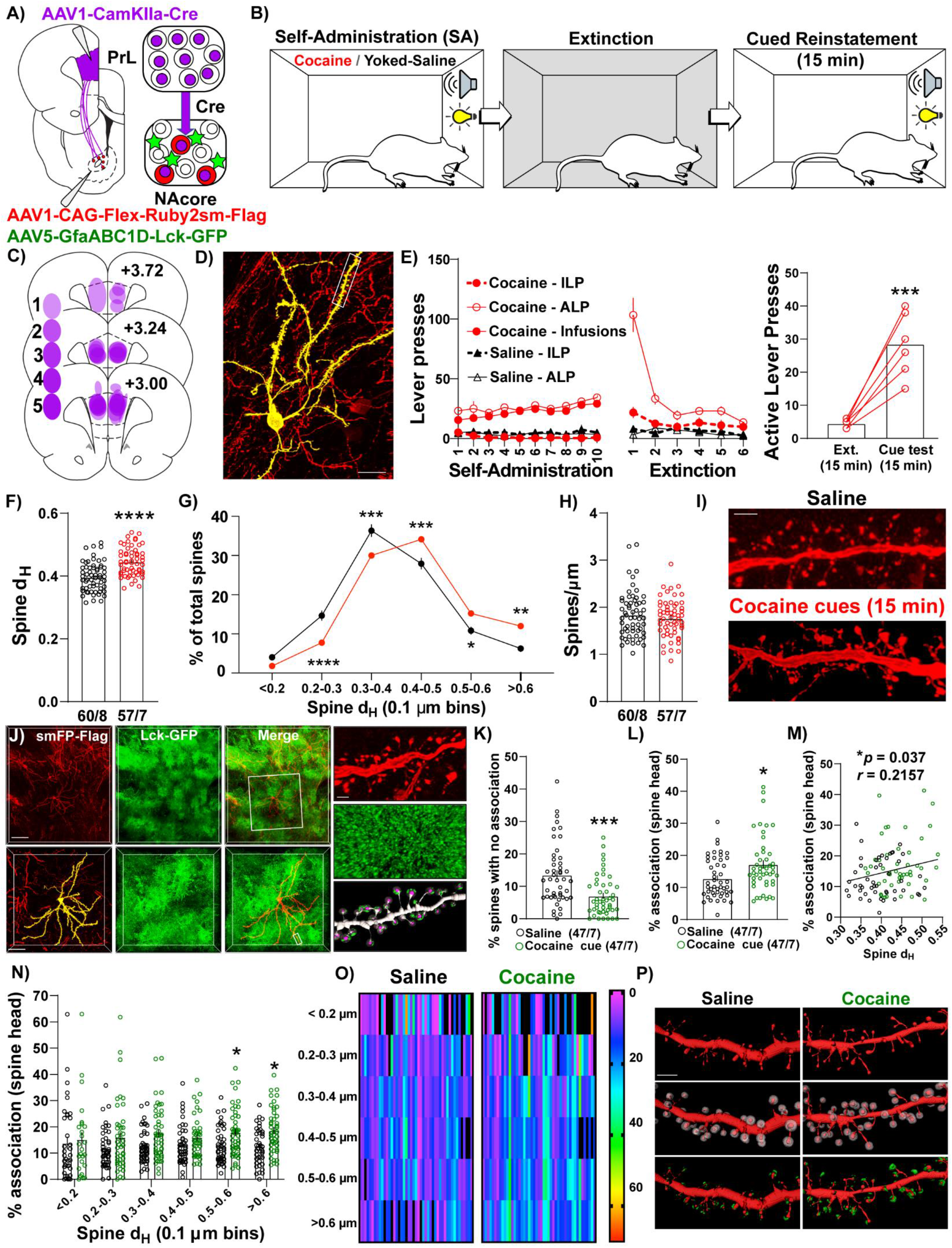
NAcore^PrL^ neurons undergo the structural component of transient potentiation which is associated with enhanced astrocyte association at spines during cue-induced reinstatement. **A)** Surgical schematic for labeling NAcore^PrL^ neurons for dendritic spine morphology and astrocyte association imaging. **B)** Behavioral timeline. One cocaine animal was removed for off-target virus expression. Animals either underwent cocaine SA (*n*=7) or received yoked saline infusions (*n*=8), extinction, and cocaine SA animals underwent cue-induced reinstatement for 15 minutes. **C)** Heat map of viral spread in the PrL cortex revealed by Cre immunohistochemistry. **D)** Representative Flex-smFP labeled NAcore^PrL^ MSN in the NAcore. **E)** ILP, ALP, and infusions earned for cocaine SA and yoked saline rats during SA and extinction (left) and active lever presses during the 15 minutes of cue-induced reinstatement compared to the first 15 minutes of the last extinction session (right); \*\*\**p*<0.001 compared to extinction. **F)** Dendritic spine head diameter (d_H_) was increased at 15 minutes of reinstatement in cocaine SA animals compared to yoked saline animals sacrificed at an extinction baseline (Yoked saline – 60 dendrite segments from 8 animals; cocaine – 57 segments from 7 animals); \*\*\*\**p*<0.0001 compared to yoked saline. **G)** Frequency distribution of binned dendritic spine d_H_ showing a general rightward shift in the distribution in cocaine reinstating animals. \**p*<0.05, \*\**p*<0.01,\*\*\**p*<0.001, \*\*\*\**p*<0.0001 compared to yoked saline. **H)** There was no difference in dendritic spine density between groups. **I)** Representative dendritic spine segments from yoked saline and cocaine cue reinstating rats. Scale bar=2µm. **J)** Representative image of Flag-tagged Ruby2sm-labeled NAcore^PrL^ MSN and surrounding Lck-GFP-labeled astrocytes (top, scale bar=100µm) and high magnification inset (bottom, scale bar=50µm) of boxed region in top panel. Right-High magnification image of boxed region in bottom left panel showing a dendritic spine segment (top, scale bar=2µm), Lck-GFP (middle), and Lck-GFP (green) confined to the region surrounding each dendritic spine head (purple) across the dendrite. **K-L)** For astrocyte interaction analyses, one yoked saline animal was removed from association analyses due to weak Lck-GFP virus expression relative to all other animals (*n*=8 segments). An additional 5 segments were removed from the yoked saline group, and 10 were removed from the cocaine group, due to either uneven Lck-GFP intensity across the dendrite or because the imaged astrocyte only ensheathed ≤50% of the dendrite. Cue-induced reinstatement of cocaine seeking decreases the percent of dendritic spines that have no association (K), while increasing the average percent association across the dendrite (L). \**p*<0.05, \*\*\**p*<0.001 compared to yoked saline. **M)** The percent association positively correlated with the average dendritic spine d_H_. **N)** Percent association as a function of spine d_H_ separated by spine size bin. \**p*<0.05 compared to yoked saline. **O)** Heat map of data in N. Warmer colors indicate greater percent association. In this instance, black refers to either 0 percent association or a bin with no spines. Each dendrite sampled is represented as 1 column. **P)** Representative NAcore^PrL^ dendrites sampled (top), the same dendrite with expanded region surrounding each spine (middle), and astrocyte membrane physical volume confined to the region surrounding each spine head (bottom). F-H, K-O – each data point represents the average value for an individual dendrite.

Next, we evaluated whether astrocyte PAPs undergo morphological plasticity at dendritic spine heads of NAcore^PrL^ neurons during seeking and whether this occurs at specific subsets of spines (Figure 4J, S1A). Interestingly, compared to yoked-saline controls, cue-induced cocaine seeking animals had a significantly lower percent of spines without any astroglial association (Welch’s-corrected two-tailed t-test, t(80.07)=3.933, *p*=0.0002, Figure 4K), and the average astroglial association with dendritic spine heads on NAcore^PrL^ neurons was significantly increased during cocaine seeking (Mann-Whitney test, U=781, *p*=0.014, Figure 4L), which positively correlated with the dendrites’ average spine d_H_ (*r*=0.216, *p*=0.037, Figure 4M). We also examined which NAcore^PrL^ spines showed the increased astroglial association. When analyzing the astroglial association across binned spine head sizes (Figure 4N-P), we observed a main effect of cue (F(1,92)=7.08, *p*=0.009) and an increase in the percent astrocyte association with the largest NAcore^PrL^ spines (Bonferroni post-hoc, 0.5-0.6 µm–*p*=0.049, >0.6 µm–*p*=0.023). Importantly, these effects were not due to differences in the intensity of GFP in the astrocyte processes (Figure S1B).

We also asked whether these adaptations occur 24 hours after the last SA session given that repeated cocaine increases basal dopamine concentrations in the NAc 24 hours after the final injection (35) without altering dendritic spine density on distal dendrites of NAcore MSNs (36), and astrocytes respond to synaptically released dopamine by increasing Ca^2+^ mobility to drive gliotransmission (37), a phenomenon often paired with increased PAPs association at synapses (38). Akin to previous findings regarding spine density 24 hours after repeated daily cocaine injections (36), dendritic spine density or head diameter of NAcore^PrL^ neurons was unaltered 24 hours after cocaine SA and no changes in astrocyte-dendritic spine association were observed (Figure S5A-G).

## Discussion

Here we showed that PrL^NAcore^ neurons are required for cue-induced cocaine seeking and associated glutamate release in NAcore. Moreover, we found that a small subpopulation of NAcore^PrL^ neurons (recruited by input and density-dependent AAV1 transsynaptic transduction from the PrL cortex) are necessary for cue-induced cocaine, but not sucrose, seeking. Importantly, these neurons undergo the structural component of transient synaptic potentiation during reinstatement, as indicated by increased dendritic spine d_H_ and enhanced astrocyte association with large spines at the peak of responding for cocaine cues. These data demonstrate that NAcore^PrL^ neurons represent a crucial subset of neurons linked to cocaine, and not sucrose, seeking.

### Role of PrL^NAcore^ neurons in cocaine seeking and NAcore glutamate release

Previous findings have indicated a clear role for PrL^NAcore^ neurons in cued cocaine, but not food, seeking and the associated glutamate release (7, 15, 23). However, the contribution of PrL-dependent glutamate release in the NAcore has yet to be directly compared between cocaine and sucrose reinstating animals. Our study is the first to demonstrate in real time that chemogenetic inhibition of PrL^NAcore^ neurons suppresses both cue-induced seeking *and* the associated elevations in NAcore glutamate release. Interestingly, chemogenetic inhibition of PrL^NAcore^neurons normalized cue-induced glutamate release in the NAcore to levels that were nearly identical to those observed in animals undergoing sucrose seeking. Thus, PrL^NAcore^ neurons represent the primary source of cue-induced glutamate release in the NAcore to drive cocaine seeking.

### Identification and characterization of NAcore^PrL^ Neurons

Recent advances in AAV1-mediated transsynaptic transduction allow for the manipulation of neurons by virtue of their inputs (31, 32). Importantly, transsynaptic transgene expression is highly dependent on the number of active connections made between the upstream and downstream neuron (32). Consistent with this property of AAV1 transsynaptic transduction, we found that 100% of virally transduced NAcore^PrL^ neurons responded when optically stimulating PrL terminals expressing DIO-hChR2, yet 58% of non-transduced cells either did not respond (33%) or showed a negligible response (≤10 pA, 25%), and EPSC amplitude was significantly greater in NAcore^PrL^ neurons when compared to neighboring unlabeled cells. We observed that NAcore^PrL^ neurons represent ∼10% of total NeuN^+^ neurons in the NAcore, the majority of which were MSNs. While this estimate is likely conservative given that this relies upon 100% transduction in the PrL, 1:1 transsynaptic transfer and 100% transduction in the NAcore, our data do support the existence of a subpopulation of NAcore neurons receiving the highest density and functionally relevant PrL input.

### NAcore^PrL^ neurons are required for cue-induced reinstatement of cocaine, but not sucrose, seeking

The NAcore is a critical region mediating cue-induced seeking for all drugs of abuse (39-41), whereby drug-induced cellular adaptations distinct from those induced by natural rewards set the stage for relapse vulnerability. Given the importance of the corticostriatal pathway in drug seeking (Figure 1) and the heterogeneity of PrL input density and functional innervation of NAcore neurons(18, 22), we used AAV1 vectors to isolate and manipulate the NAcore MSNs most heavily innervated by the PrL cortex. We found that chemogenetic inhibition of NAcore^PrL^ neurons suppressed cue-induced cocaine, but not sucrose, seeking. One limitation of our approach is that NAcore^PrL^ neurons were transduced by virtue of their input from the PrL and not by virtue of their input *and* cell type. This is important given that we observed NAcore^PrL^ neurons constitute a mix of D1 and D2-expressing MSNs as well as interneurons. We suspect that chemogenetic inhibition of D1-NAcore^PrL^ and D2-NAcore^PrL^ neurons would have dichotomous effects on cued seeking, consistent with previous reports (42, 43). Our AAV1-derived estimates indicate that only 20% of total D2-MSNs receive dense PrL input, with NAcore^PrL^ neurons being approximately 50% D2-MSNs and by estimation 36% D1-MSNs. While just 7% of NAcore^PrL^ neurons were PV interneurons, 60% of total PV interneurons were labeled with our AAV1 transsynaptic vector, indicating that a large portion of total PV interneurons receive this functionally relevant PrL cortical input. As such, future experimentation will need to address whether chemogenetic inhibition of the population of PrL input receiving PV interneurons is sufficient to suppress cue-induced reinstatement. Indeed, suppressing PV interneuron activity in the NAc reduces amphetamine sensitization in mice (44). Overall, the interaction between NAcore^PrL^ subtypes in regulating cocaine-seeking behavior is undoubtedly complex, and future studies will need to be performed to delineate the involvement of each specific subtype.

Regardless of the cell types involved, activity in NAcore^PrL^ neurons appears to be critical for overall NAcore activation during cocaine seeking given that chemogenetic inhibition of NAcore^PrL^ neurons led to an overall suppression of Fos^+^ neurons in the NAcore. Consistent with recent reports (45), we observed that overall levels of Fos induction in the NAcore per μm^3^ were similar in cocaine and sucrose reinstating mCherry control animals. However, Fos induction was significantly higher in NAcore^PrL^ neurons in mCherry cocaine animals compared to mCherry sucrose. Further, the magnitude of Fos activation in NAcore^PrL^ neurons positively correlated with the amount of seeking in cocaine, but not sucrose, animals. Collectively, these data demonstrate that NAcore^PrL^ neuronal activity is necessary for cue-induced cocaine, but not sucrose, seeking. Given that inhibition of PrL^NAcore^ neurons during cocaine seeking normalized NAcore glutamate levels to that of sucrose seeking animals (Figure 1), we speculate that both potentiated PrL-dependent excitatory transmission in the NAcore combined with cellular adaptations in NAcore^PrL^ neurons following extinction from cocaine, but not sucrose, SA sets the stage for relapse vulnerability.

### NAcore^PrL^ neurons and astrocytes undergo morphological plasticity during cocaine seeking

Cue-induced seeking for multiple drugs of abuse elicits time-dependent increases in synaptic strength within the NAcore MSNs, including increased AMPA/NMDA ratio and enlarged dendritic spine d_H_ (46), the degree of which positively correlates with drug-seeking behavior (23). As a recent addition to established cue-induced neuronal structural plasticity, heroin seeking has been linked to increased astrocyte association with NAcore synapses. This cue-induced astrocytic structural plasticity limits drug seeking (*i*.*e*. promotes within-session extinction), likely by increasing glutamate clearance (26). However, previous studies did not provide evidence that increased astrocyte association occurred at relevant synapses. Our data are the first to demonstrate that NAcore^PrL^ MSNs undergo increased dendritic spine d_H_ during the peak of reinstatement, as well as increased association of perisynaptic astrocyte processes with the most potentiated spines on NAcore^PrL^ neurons, an effect that was not observed 24 hours following cocaine SA. These data suggest that cue-induced motility of astroglial processes and increased interaction with structurally potentiated synapses at NAcore^PrL^ MSNs serves to limit glutamate spillover during cocaine seeking.

In conclusion, we demonstrate that PrL^NAcore^ neurons are necessary for cue-induced glutamate release in the NAcore and cocaine seeking. Moreover, a subset of NAcore neurons receiving dense and functional PrL cortical input are required for cocaine and not sucrose seeking, and undergo transient cue-induced morphological plasticity, including dendritic spine expansion and astrocyte association at potentiated spines. Taken together, NAcore^PrL^ neurons represent a novel target for understanding how drug-induced adaptations in functional circuits embedded within a known network serve as the molecular basis for craving and relapse vulnerability.

## Acknowledgments

This work was funded by DA054154 (MDS), DA050427 (BMS), DA051650 (JMO), DA032708 (CWC), DA047845 (SMB) and NIDA Center on Opioid and Cocaine Addiction (DA046373).

## Supplemental information

### Materials and methods

#### Animal subjects and surgery

Male Sprague-Dawley rats were purchased from Charles Rivers Laboratories (Wilmington, MA,∼275-300 g). Rats were single-housed in a temperature and humidity controlled vivarium with a 12:12 hour reverse light/dark cycle (lights off at 6AM). All experiments were conducted during the dark phase. Animals were allowed to acclimate for 5 days prior to undergoing experiments with food and water available *ad libitum*. Prior to surgery, animals were anesthetized with a ketamine (66 mg/kg, i.p.) and xylazine (1.33 mg/kg) cocktail and received ketorolac (Sigma, 2 mg/kg i.p.) for analgesia. Animals undergoing intravenous (i.v.) cocaine (or yoked-saline) SA were implanted with a chronic indwelling catheter as previously described (12, 47). Catheters exited via a small incision in the shoulder blades. Following surgery, catheters were flushed with Cefazolin to prevent bacterial infection. Catheters were flushed daily for 5 days post-surgery with 50 µl of taurolidine citrate catheter lock solution (Access technologies) to maintain patency. Animals were then secured in a stereotaxic apparatus (David Kopf instruments, model 942). Rats then received an intra-prelimbic (PrL) cortical (AP: + 2.8mm, ML: +/-0.6mm, DV: −3.8 mm relative to bregma) virus injection and an intra-nucleus accumbens core (NAcore, AP: +1.7mm, ML: +/-1.6mm, DV:-7mm relative to bregma) virus injection (see table 1 for viral vector information). Injections were performed over a period of 5 minutes using a Nanoject II (0.75 µl/ hemisphere, Drummond Scientific, Broomall, PA). Injectors were left in place for 5 minutes to facilitate diffusion away from the injection site, then slowly retracted. The incision was then sutured closed and animals were allowed to recover from anesthesia.

#### Cocaine and sucrose SA

Cocaine and sucrose SA (SA) were performed as previously described (12, 30, 47, 48). Briefly, rats were mildly food-deprived (∼15 g rat chow/day) to increase exploration prior to beginning SA and were then maintained at 20 g/day for the remainder of the experiment. Rats were weighed daily, and catheters were flushed with sterile saline prior to daily SA sessions. Rats then underwent SA (Fixed ratio (FR) 1 schedule of reinforcement) in standard MedPC operant chambers (Med Associates, St. Albans, VT) fixed with two retractable levers, a house light, tone generator, sucrose dispenser, and two lights above each lever) for two hours/day (12-14 days). Presses on the active lever elicited a light and tone cue-complex followed by a single infusion of cocaine or a single 45 mg flavored sucrose pellet (Bioserv, Flemington, NJ, #F0025) followed by a 20 second timeout period. Presses on the inactive lever had no programmed consequence. The day after the final SA session, rats either entered extinction training or received an intra-NAcore cannula implantation for electrochemical experiments (see below).

#### Perfusions and immunohistochemistry

Immunohistochemistry was performed as previously described (8, 30, 49). Briefly, rats were transcardially perfused with 5% buffered formalin (150 ml) following a pre-flush with 0.1M phosphate buffered saline (PBS, 120 ml) at a rate of 60 ml/minute. For dendritic spine morphology and astrocyte association at dendritic spine experiments, rats were pre-flushed with 0.1M phosphate buffer (PB, 120 ml) followed by a transcardial perfusion with 4% granular paraformaldehyde (180 ml) in 0.1M PB. Brains were extracted and post-fixed in the same fixative for 24 hours. A vibrating microtome (Leica) was used to section brains. Free-floating coronal sections containing the PrL cortex (60 µm) and NAcore (60-80 µm) were blocked in 0.1M PBS with 2% Triton X-100 (PBST) with 2% normal goat serum (NGS, Jackson Immuno Research, Westgrove, PA) for 2 hours at room temperature with agitation. Sections were then incubated overnight at 4°C with agitation in the appropriate primary antisera diluted in 2% PBST with 2% NGS (see Table 2), washed 3 times for 5 minutes in PBST, then incubated in the appropriate secondary antisera (see Table 2) diluted in PBST with 2% NGS for 2-5 hours at room temperature with agitation. All secondary antisera were raised in goat, conjugated to Alexa fluorophores, were used at a concentration of 1:1000, and were purchased from Invitrogen (Carlsbad, CA). Sections were then washed 3 times for 5 minutes in PBST, mounted on SuperFrost+ slides, and cover slipped with ProLong™ Gold Antifade. Slides were stored at 4°C protected from light until imaging.

### Microscopy

#### Mapping of virus expression and, colocalization of mCherry/Flag with different cell types, and Fos

All imaging was performed by an investigator blind to experimental conditions. To determine accuracy of microinjections in both the PrL cortex and NAcore, a Leica THUNDER Imager Tissue equipped with 488, Cy3, and Ct5 filter cubes (Leica microsystems) was used. Exposure time was held relatively constant between images. Virus expression was then mapped by an investigator blind to experimental conditions, and expression profiles were overlaid to generate a heatmap.

AAV1 vectors expressing Cre recombinase are known to have transsynaptic properties whereby transduction of neurons, and thus expression of Cre, downstream from the injection site is known to occur in a manner dependent on vesicular release as well as the strength of the connection between the neurons (31, 32). To determine what cell types in the NAcore are transduced by AAV-CamKIIa-Cre when injected into the PrL cortex, tissue from Yoked Saline controls, animals undergoing 15 minutes of reinstatement, and animals undergoing 2 hours of cue-induced reinstatement were immunohistochemically processed for various markers and a Leica SP8 laser-scanning confocal microscope was used to image virally-transduced neurons in the NAcore in addition to either NeuN, pre-pro Enkephalin (ppENK), nNOS, Parvalbumin (PV), or Choline acetyltransferase (ChAT) in separate IHC runs. mCherry/Flag were imaged using an OPSL 552nm laser line whereas all other markers were imaged using a Diode 638nm laser line. Pinhole size, laser power, gain, and voxel size were held constant between images within each experimental IHC run and images were acquired with either a 10X or 20X air objective, but the same objective was used throughout the experimental run for each marker.

For detection of Fos, virally-transduced neurons (mCherry^+^) and Fos^+^ nuclei in the NAcore from animals undergoing cue-induced reinstatement were imaged with a 10X objective with 2X digital zoom. Care was taken to ensure that the field imaged contained mCherry^+^ neurons within each quadrant and that images were only acquired in the NAcore. As before, all parameters were held constant between images.

#### Imaris analyses

##### Colocalization of mCherry/Flag with different cell types and Fos activated neurons

All Imaris image analyses were performed on 3D reconstructed Z-stacks by an investigator blind to experimental conditions. Non-deconvolved images were exported to Imaris (v 9.0, Bitplane) for analyses. For all images, a constant baseline subtraction was applied to the non-mCherry/Flag channel in order to apply a threshold cutoff. Next, the spot tool in Imaris was used to semi-manually label mCherry^+^/Flag^+^ cells as well as NeuN^+^, nNOS^+^, ppENK^+^, PV^+^, ChAT^+^, or Fos^+^ cells/nuclei. Due to the low population of interneurons, the number of mCherry^+^/interneuron^+^ neurons were manually counted and normalized to the total number of mCherry^+^ cells or the total number of interneurons in the image to generate a percentage of total mCherry^+^ cells that were nNOS^+^, PV^+^, or ChAT^+^ as well as the inverse (percentage of interneurons that were transduced). For ppENK and NeuN, iterative processing and analyses were performed whereby spots were built on mCherry^+^/Flag^+^ cells, then the ppENK or NeuN signal contained within the transduced cells was masked, and spots were built on the new channel as well as the original channel. For Fos, the colocalize spots MatLab extension (5 µm threshold) was used after spots were built on mCherry^+^ cells and Fos^+^ nuclei. Exported variables included the number of mCherry^+^/Flag^+^ cells,the number of co-labeled (of Fos^+^) cells, the percentage of mCherry^+^/Flag^+^ cells that were co-labeled, as well as the percentage of NeuN^+^, nNOS^+^, ppENK^+^, PV^+^, or ChAT^+^ cells that were virally-transduced. The number of Fos^+^ neurons was normalized to the dataset volume (in µm^3^). For Fos analyses, data was collapsed across hemispheres, then sections, and expressed as an animal average.

**Figure S1.**
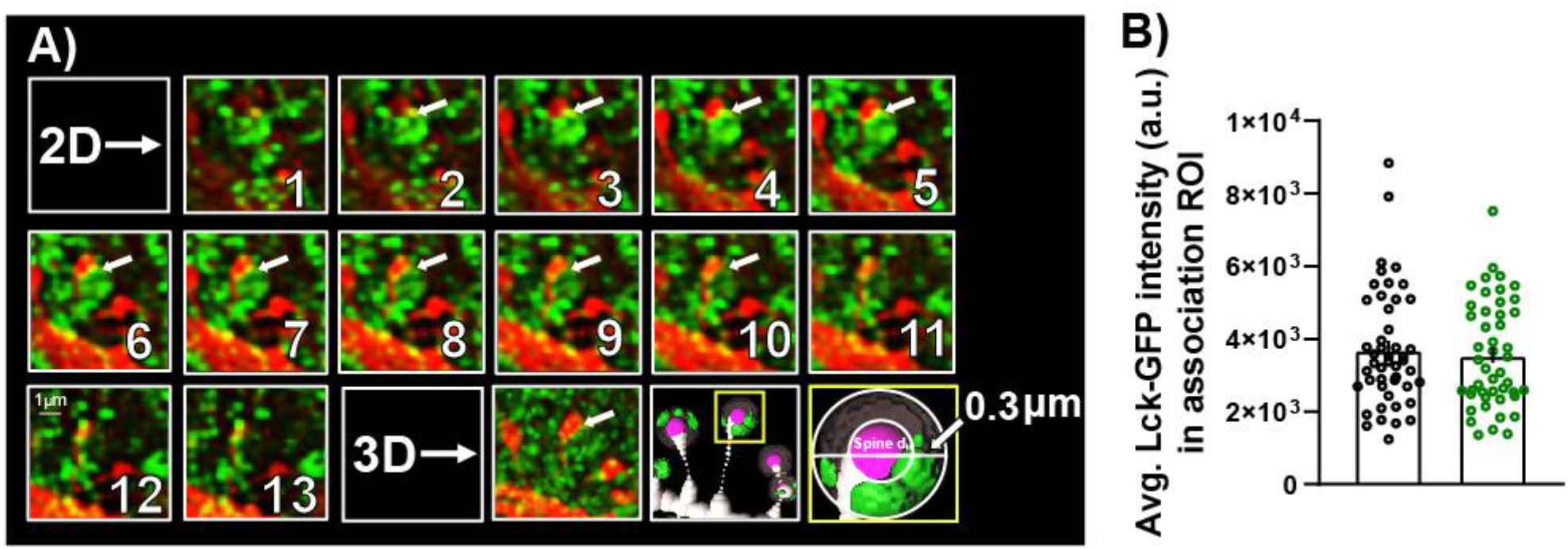
Z-series of astrocyte association at dendritic spines and intensity control data, pertaining to Figure 4. A) Individual Z-steps showing an individual spine (marked with an arrow) with Lck-GFP expression surrounding the spine at 13 different focal planes, as well as a 3D render of the same spine. Bottom right – Surface rendering of the dendritic spine and astrocyte membrane surrounding the spine with the spine head rendered as a sphere. Grey area indicates the expanded spine head sphere in 300 nm in all directions. Green render indicates astrocyte in association region of interest (ROI). B) Average Lck-GFP intensity in the association ROI for each dendrite segment analyzed for yoked saline and cocaine cue reinstating animals. There was no difference between groups (t(92)=0.482, *p*=0.631).

**Figure S2.**
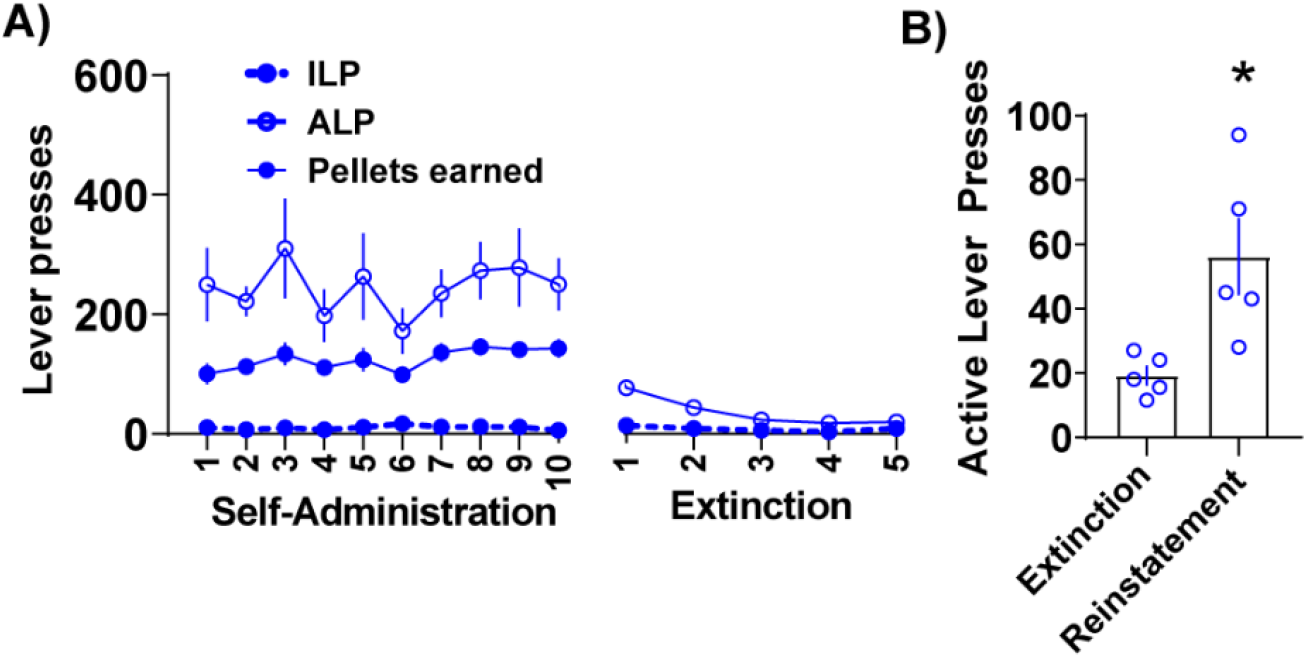
Behavior data pertaining to Figure 1. A-B) Sucrose SA and extinction (A) and reinstatement (B) lever pressing for animals undergoing electrochemical recordings (*n*=5) during reinstatement. \**p*<0.05 compared to extinction.

**Figure S3.**
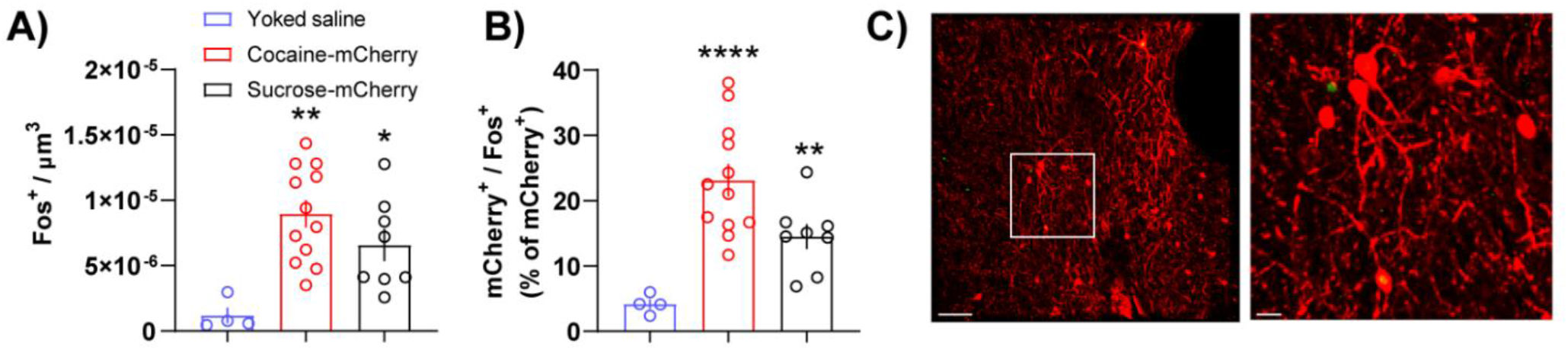
Cue-induced Fos is increased in cocaine and sucrose animals compared to yoked saline control animals. A) Compared to yoked saline controls (*n*=4, from dendritic spine and astrocyte association experiment – Figure 4), cocaine and sucrose animals expressing mCherry control in NAcore^PrL^ neurons show increased Fos in the NAcore (One-way ANOVA: F(2,21)=8.126, *p*=0.0024, Dunnett’s post hoc, *p*<0.05). B) Fos was also increased in NAcore^PrL^ neurons in the two cocaine groups compared to yoked saline (Brown-Forsythe-corrected ANOVA: F(2,18.92)=20.14, *p*<0.0001, Dunnett’s T3 multiple comparison, *p*<0.01). C) Representative Fos expression from a yoked saline control animal. Scale bars = 50µm (left) and 20 µm (right).

**Figure S4.**
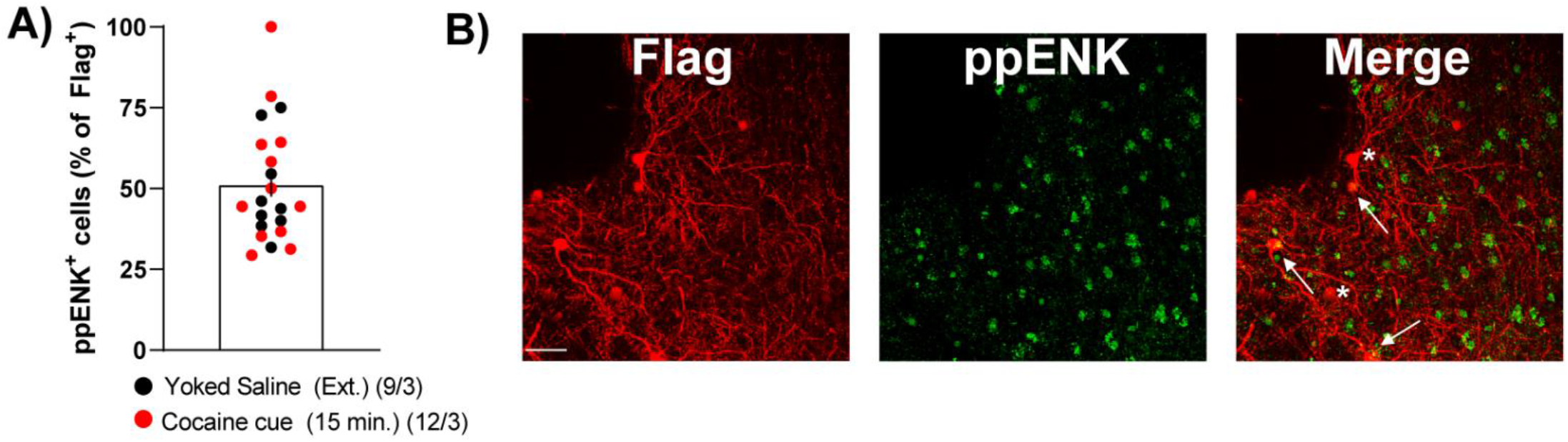
Estimates of NAcore^PrL^ D2-expressing neuron frequency, pertaining to Figure 4. A) ∼50% of cells labeled for dendritic spine morphology measurements were ppENK^+^. There was no difference between groups in the percent of labeled neurons that were ppENK^+^ (t(19)=0.44, *p*=0.663). B) Representative Flag-tagged-Flex-smFP neurons (Red) and immunohistochemical detection of ppENK (green). Arrow heads point to Flag^+^/ppENK^+^ neurons. Asterisks indicate Flag^+^/ppENK^-^ neurons. Scale bar=50µm. Tissue was processed from animals expressing Flex-smFP undergoing yoked-saline and sacrificed 24 hours after the final extinction session (*n*=3) or 15 minutes into cued reinstatement of cocaine seeking (*n*=3).

**Figure S5.**
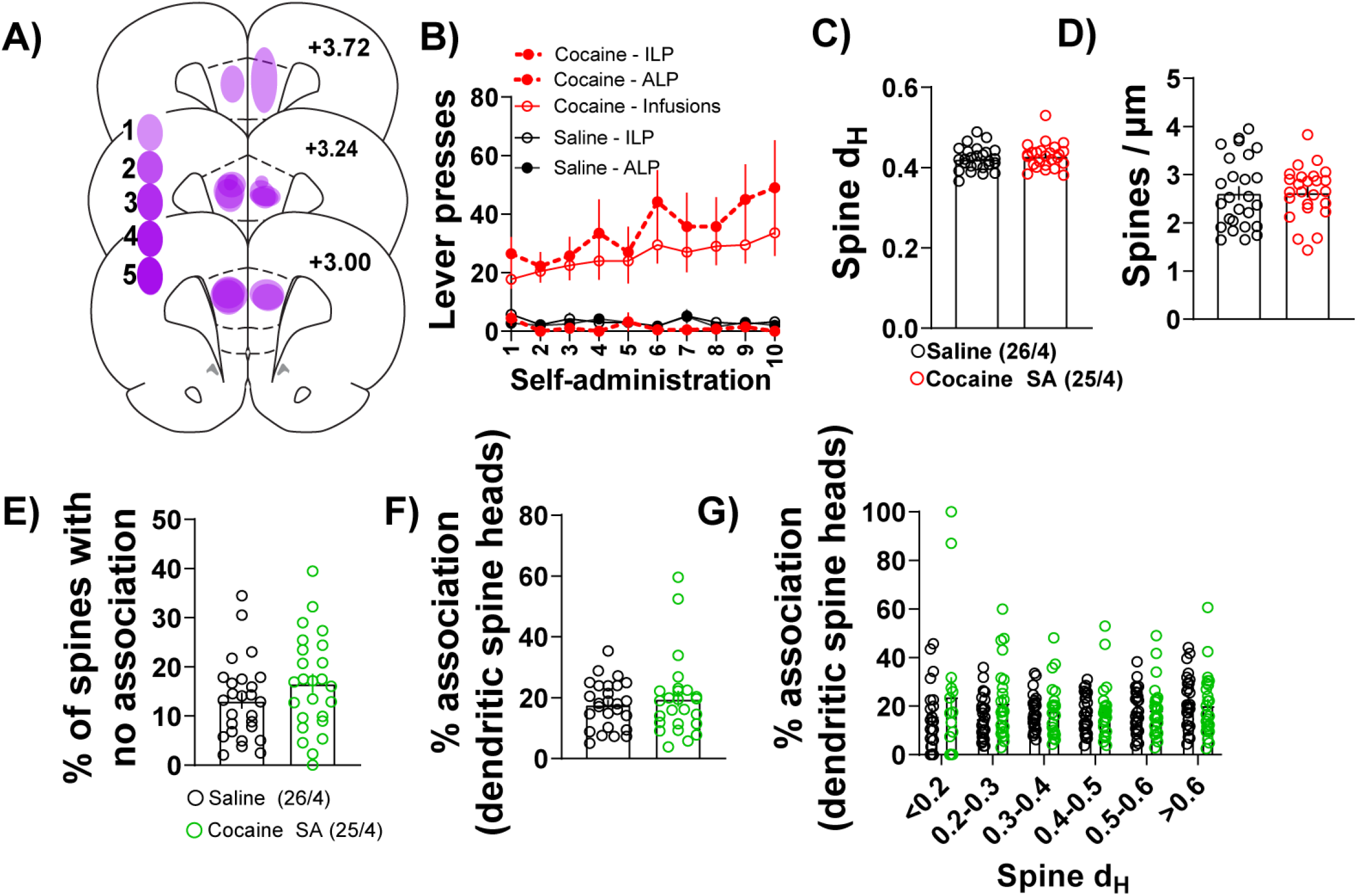
There were no changes in NAcore^PrL^ dendritic spine morphology or astrocyte association at spine heads 24 hours after the final cocaine SA session. A) Heat map of viral spread in the PrL cortex as revealed by Cre immunohistochemistry. B) Cocaine (*n*=4) and yoked saline (*n*=4) ILP, ALP, and infusions earned over the last 10 days of SA. C-D) There was no difference in spine d_H_ (C - unpaired t-test, t(49)=0.834, *p*=0.408) or (D – unpaired t-test, t(49)=0.101, *p*=0.92) dendritic spine density on NAcore^PrL^ neurons 24 hours after the final SA session. E-G) There was no difference in the percent of spines per dendrite that showed no association (E – unpaired t-test, t(49)=1.392, *p*=0.17) nor the average association at dendritic spine heads (F – Welch’s-corrected t-test, t(38.78)=0.647, *p*=0.522), and there was no specific dendritic spine d_H_ bins that showed any alterations in percent association at spine heads (G, significant spine d_H_ bin by treatment interaction, (F(5,229)=2.768, *p*=0.02, no significant Bonferroni post-hoc test).

**Table S1.** Non-significant behavioral data.

| Figure | Test | Comparison | Statistics |
| --- | --- | --- | --- |
| 1C | Two-way RM ANOVA | hM4Di vs. mCherry, ALPs during SA | Non-significant session by group interaction (F(9,189)=0.971, $p=0.465$) |
| 1C | Two-way RM ANOVA | hM4Di vs. mCherry Infusions earned during SA | Non-significant session by group interaction (F(9,189)=1.320, $p=0.229$) |
| 1E | Two-way RM ANOVA | hM4Di vs. mCherry ALPs across time | Non-significant session by group interaction (F(7,147)=0.798, $p=0.599$) |
| 3E | Two-way RM ANOVA | hM4Di vs. mCherry ALPs during SA | Non-significant session by group interaction (F(9,189)=1.089, $p=0.372$) |
| 3E | Two-way RM ANOVA | hM4Di vs. mCherry Infusions earned during SA | Non-significant session by group interaction (F(9,189)=0.761, $p=0.65$) |
| 3E | Mixed-effects model | hM4Di vs. mCherry ALPs during extinction | Non-significant session by group interaction: F(6,101)=0.423, $p=0.863$) |
| 3F | Two-way RM ANOVA | hM4Di vs. mCherry ILPs during extinction and reinstatement | Non-significant group by test interaction (F(1,21)=0.471, $p=0.5$) |
| 3H | Two-way RM ANOVA | hM4Di vs. mCherry ALPs during SA | Non-significant session by group interaction (F(9,126)=1.278, $p=0.255$) |
| 3H | Two-way RM ANOVA | hM4Di vs. mCherry pellets earned during SA | Non-significant session by group interaction: (F(9,126)=1.749, $p=0.085$) |
| 3H | Mixed-effects model | hM4Di vs. mCherry ALPs during extinction | Non-significant session by group interaction (F(5,65)=0.394, $p=0.851$) |
| 3I | Two-way RM ANOVA | hM4Di vs. mCherry ILPs during extinction and reinstatement | Non-significant session by group interaction (F(1,14)=3.452, $p=0.084$) |

#### Key resource table

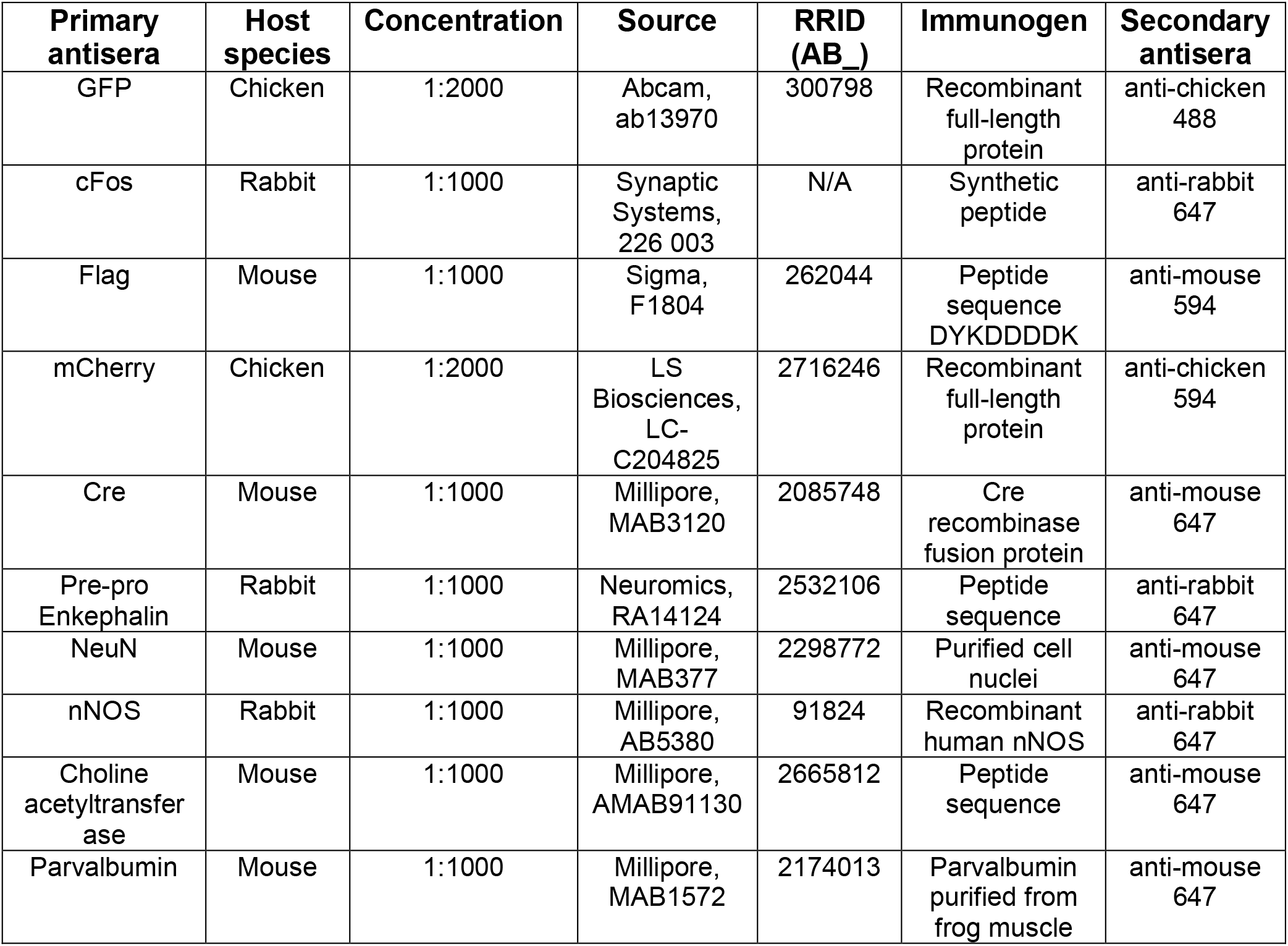

| **Primary antisera** | **Host species** | **Concentration** | **Source** | **RRID (AB_)** | **Immunogen** | **Secondary antisera** |
| --- | --- | --- | --- | --- | --- | --- |
| GFP | Chicken | 1:2000 | Abcam, ab13970 | 300798 | Recombinant full-length protein | anti-chicken 488 |
| cFos | Rabbit | 1:1000 | Synaptic Systems, 226 003 | N/A | Synthetic peptide | anti-rabbit 647 |
| Flag | Mouse | 1:1000 | Sigma, F1804 | 262044 | Peptide sequence DYKDDDDK | anti-mouse 594 |
| mCherry | Chicken | 1:2000 | LS Biosciences, LC-C204825 | 2716246 | Recombinant full-length protein | anti-chicken 594 |
| Cre | Mouse | 1:1000 | Millipore, MAB3120 | 2085748 | Cre recombinase fusion protein | anti-mouse 647 |
| Pre-pro Enkephalin | Rabbit | 1:1000 | Neuromics, RA14124 | 2532106 | Peptide sequence | anti-rabbit 647 |
| NeuN | Mouse | 1:1000 | Millipore, MAB377 | 2298772 | Purified cell nuclei | anti-mouse 647 |
| nNOS | Rabbit | 1:1000 | Millipore, AB5380 | 91824 | Recombinant human nNOS | anti-rabbit 647 |
| Choline acetyltransferase | Mouse | 1:1000 | Millipore, AMAB91130 | 2665812 | Peptide sequence | anti-mouse 647 |
| Parvalbumin | Mouse | 1:1000 | Millipore, MAB1572 | 2174013 | Parvalbumin purified from frog muscle | anti-mouse 647 |

